# Mechanochemical modeling of exercise-induced skeletal muscle hypertrophy

**DOI:** 10.64898/2025.12.17.694686

**Authors:** Ingvild S. Devold, Marie E. Rognes, Padmini Rangamani

## Abstract

Skeletal muscle displays remarkable plasticity, adapting its size and strength in response to mechanical loading, particularly, from exercise. This process, known as hypertrophy, is fundamental to athletic training and rehabilitation, but is challenging to quantitatively predict due to its multifactorial, multiscale nature. Specifically, skeletal muscle hypertrophy results from an integration of macroscopic mechanical stimuli with the intracellular signaling pathways that govern muscle growth. In this work, we present a multiscale computational model that mechanistically integrates these mechanical and biochemical stimuli and offers a framework for predicting the outcomes of different types of exercise on skeletal muscle growth. The framework couples a transversely isotropic hyperelastic model for tissue-level mechanics with a system of ordinary differential equations representing the IGF1-AKT-mTOR-FOXO signaling pathway, a key regulator of protein synthesis and degradation. We link these scales using a volumetric growth model, where the signaling dynamics inform a growth tensor that drives changes in muscle cross-sectional area. This approach enables the simulation of long-term muscle adaptation, providing a mechanistic tool to investigate how different exercise protocols lead to macroscopic hypertrophy. Simulations from our model capture the temporal dynamics of hypertrophy under varying load protocols and highlight how feedback between protein synthesis and muscle growth regulates the dose-response relationship to prevent unbounded growth. Using muscle geometries derived from the Visible Human dataset, we study how human variations in muscle geometry affect hypertrophy. Finally, we demonstrate that the mechanochemical coupling between muscle geometry and signaling not only predicts macroscopic shape changes but also provides buffering from local signaling heterogeneity. Ultimately, this framework offers a predictive computational tool for optimizing training regimens and understanding the multiscale determinants of muscle adaptations.

## Introduction

Skeletal muscle is remarkably plastic, constantly adapting its size and structure to mechanical stimuli resulting from different movements. Such plasticity is not only critical for muscle’s ability to respond to the physical demands placed on it^1, 2^ but is also critical to athletic training, rehabilitation, and overall health. Different types of exercise such as high-intensity sprints, high-load resistance training, or long-duration endurance training, induce distinct patterns of muscle activation that are characterized by variations in duration, amplitude, and frequency (Figure 1A). While the effects of exercise on skeletal muscle function are empirically well-documented^3^, the underlying connections between molecular mechanisms and mechanical responses driving these adaptations remain poorly understood^4^. This is largely because they represent a profoundly multiscale phenomenon, which require an integration of tissue-level mechanics and subcellular biochemical signal transduction that has proven challenging to model.

**Figure 1.**
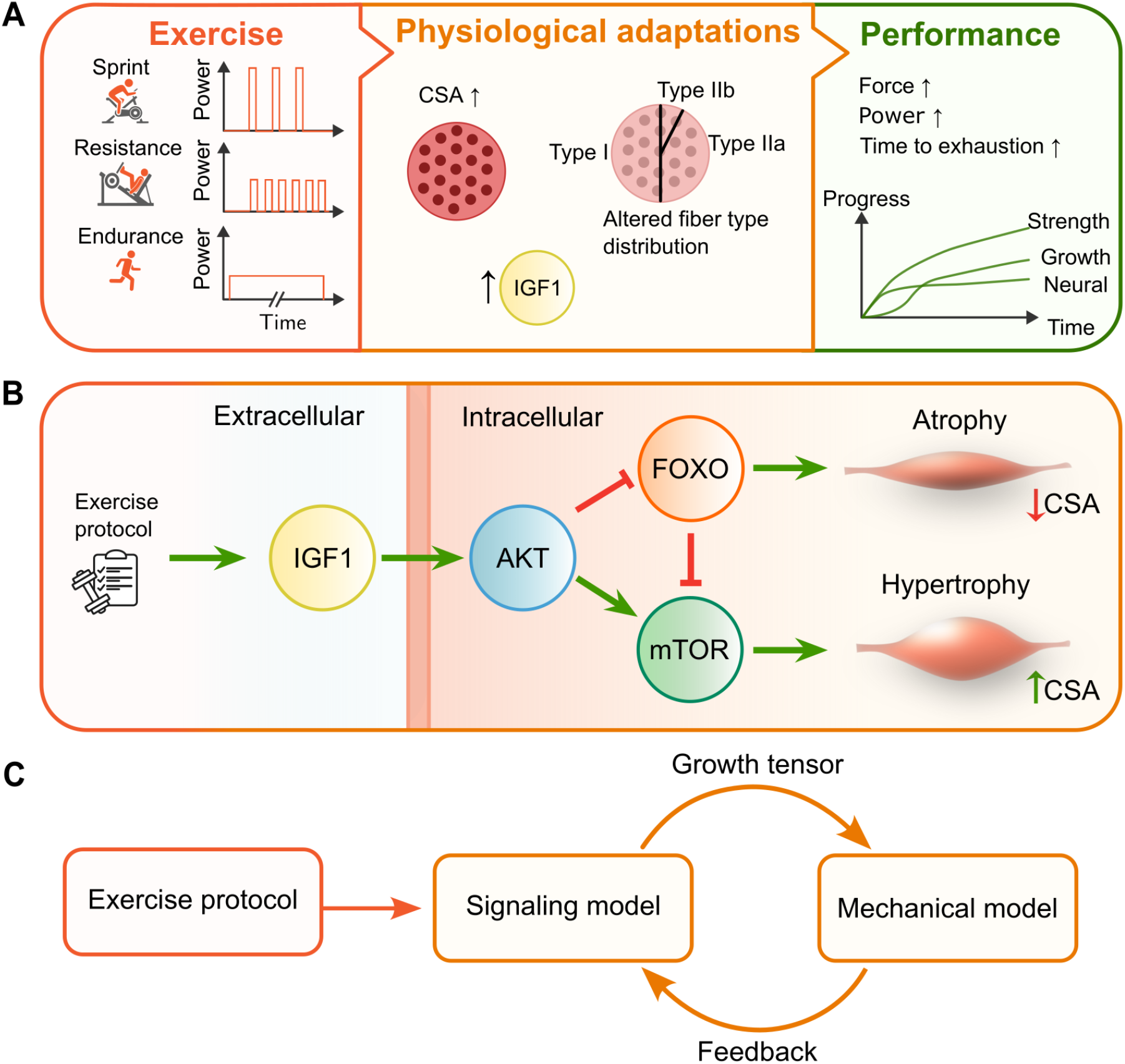
Conceptual overview of exercise-induced muscle adaptation and the modeling framework. (**A**) Different exercise types are characterized by distinct patterns of muscle activation, leading to physiological adaptations and ultimately affecting muscle performance. (**B**) In this work, we focus on the IGF1-AKT signaling pathway, which is a key regulator of muscle hypertrophy. Mechanical stimuli through exercise activate IGF1, which in turn activates AKT. AKT promotes protein synthesis via mTOR and inhibits protein degradation via FOXO. (**C**) Schematic of the proposed multiscale model, coupling a signaling model with tissue-level mechanics to predict exercise-induced muscle growth. Abbreviations: CSA: cross-sectional area; IGF1: insulin-like growth factor 1; AKT: protein kinase B; mTOR: mechanistic target of rapamycin; FOXO: forkhead box transcription factor.

The multiscale nature of muscle spans both spatial and temporal scales^4^. Spatially, muscle tissue is organized hierarchically, from the whole organ down to fascicles, individual muscle fibers, myofibrils, and finally to the sarcomere – the fundamental 2-3 µm contractile unit composed of actin and myosin filaments^5, 6^. Temporally, muscle contraction is driven by rapid electrochemical events. This includes millisecond-timescale calcium transients triggering the ATP-fueled cross-bridge cycle, which in turn generates force and causes the relative sliding of the sarcomere filaments. Over far longer timescales, spanning weeks to months, the cumulative effects of these contractions trigger intricate signaling pathways that regulate protein synthesis and degradation^7, 8^ and changes to metabolic pathways^9^ ultimately remodeling the tissue structure. Mechanistically linking an exercise protocol to a long-term adaptation requires integrating our understanding of both muscle mechanics and cellular physiology.

From a mechanical perspective, foundational models of cellular-level force generation^10, 11^ have, with modern advances in medical imaging and computational resources, evolved into detailed three-dimensional models^12^. These models typically employ a hyperelastic material framework to accurately capture the large, non-linear deformations that muscle tissue undergoes during activity^13–15^. More recently, some models have begun to bridge scales by coupling cell-level models^16^ with tissue-level mechanics^17–19^. From a physiological perspective, there are multiple signaling pathways at play in the muscle^7^. Among these, the IGF1-AKT-mTOR-FOXO signaling pathway has been identified as a central regulator of load-induced skeletal muscle hypertrophy^20–22^. In this cascade, mechanical loading through exercise stimulates the release of insulin-like growth factor (IGF1), which activates the protein kinase B (AKT) (Figure 1B). AKT, in turn, displays a dual response: it promotes protein synthesis via mechanistic target of rapamycin (mTOR) and inhibits protein degradation by suppressing the forkhead box transcription factor (FOXO), suggesting the presence of an incoherent feedforward loop^23, 24^. The balance between these mTOR and FOXO-driven mechanisms dictates the net change in muscle protein and, consequently, muscle growth.

Computational models shed critical insights into how exercise can regulate muscle growth at different length and time scales. For short time-scale events, systems biology approaches have extensively mapped the calcium dynamics underlying excitation-contraction coupling^16, 25^, with recent work clarifying the role of store-operated Ca^2+^ entry (SOCE) in force modulation^26^, the link between mechanical loading and metabolic sensors such as AMPK^27^, and the bidirectional coupling between exercise and mitochondrial dynamics^28^. However, such signaling models typically represent the muscle tissue as a homogeneous compartment, predicting hypertrophy as a scalar quantity without accounting for the complex spatial distribution of stress and strain within the 3D tissue.

Conversely, tissue-level mechanical models excel at resolving these local deformations using hyperelastic formulations. However, they predominantly operate under the assumption of mechanical equilibrium with fixed material reference states. By treating the tissue as a static material, these models neglect the dynamic, biochemical adaptation of the tissue constituents that occurs in response to mechanical loading over time. Bridging these domains requires overcoming a fundamental timescale mismatch: instantaneous mechanical stimuli trigger signaling events that accumulate over weeks to drive morphological change, which in turn alters the mechanical environment.

The mathematical theory of volumetric growth^29–31^ offers a rigorous kinematic framework to bridge this gap. By decomposing the deformation gradient into elastic and growth contributions, this theory allows biologically driven mass accumulation from tissue growth to be coupled directly to continuum mechanics. The volumetric growth framework has previously found applications in several biological tissues^32, 33^, including cardiac^34, 35^ and brain tissue^36^. In the context of skeletal muscle, Villota-Narvaez et al.^37^ found that a multiscale mechanobiological model, which integrates the IGF1-AKT signaling pathway with continuum mechanics via a growth tensor, accurately predicts muscle hypertrophy (CSA) results under various training conditions. However, their framework focused on idealized geometries and did not fully implement the cumulative growth kinematics required to robustly track residual stresses and shape evolution over long-term training protocols.

In this paper, we seek to address the following challenges: How can we simulate growth kinematics over long timescales while properly accounting for evolving residual stresses? How can we incorporate the effect of complex, non-uniform myofiber architectures found in realistic muscle anatomies and their influence on local growth? And finally, how can we leverage advances in computational mechanics to generate a predictive framework for muscle responses to different exercise schemes? To this end, we present a multiscale computational framework that couples tissue-level mechanics with the IGF1-AKT signaling pathway (Figure 1C). We ground our approach in the mathematical theory of volumetric growth^29, 31^, which allows biologically driven mass accumulation to be coupled directly to continuum mechanics. While our work builds on the mechanobiological model of Villota-Narvaez et al.^37^, it extends the computational framework in two important directions. First, we adopt the cumulative growth framework of Goriely and Ben Amar^38^, which ensures that each growth increment is applied to a stress-free configuration, providing a robust method for handling long-term adaptation. Second, we apply this coupled model to anatomically realistic muscle geometries derived from the Visible Human dataset^39–41^, demonstrating the versatility of the approach. Using this framework, we demonstrate that high-intensity, high-frequency exercise protocols drive the most pronounced hypertrophy, reveal that complex muscle architectures result in non-uniform growth patterns, and that introducing heterogeneity in the signaling dynamics disrupts the overall growth. This integrated approach demonstrates how protocol-dependent signaling drives 3D tissue adaptation in realistic geometries, highlighting its predictive potential for both rehabilitation and peak performance sports.

## Methods

### Stress-strain relationships

Muscle tissue undergoes large deformations and exhibits a nonlinear stress-strain relationship under physiological loading. Its bundle-like structure results in a transversely isotropic material response, with distinct mechanical properties along and across the fiber directions **a**_0_^42^. Accordingly, we model the mechanical behavior of muscle using a transversely isotropic hyperelastic framework^18^. In this framework, the stress response is derived from a strain energy density function, *W*, which depends on the tissue deformation. To define this relationship, we first introduce key kinematic quantities. Let Ω ⊂ ℝ^3^ denote a fixed reference domain (open, bounded) with coordinates **X** and boundary *∂*Ω. We let **u** = **x** − **X** denote the displacement from this reference configuration to the current configuration. The deformation gradient is given by **F** = ∇_**X**_**u** + **I**, and the corresponding right Cauchy-Green deformation tensor is **C** = **F**^*T*^ **F**. The local volume change is measured by the Jacobian *J* = det(**F**). The strain energy function for a transversely isotropic material is defined through the invariants of **C**: the first invariant *I*_1_ = tr(**C**) and the fourth quasi-invariant *I*_4_ = **a**_0_ · **Ca**_0_. Finally, the fiber stretch is given by 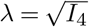.

The stress experienced by the material is quantified by the first and second Piola–Kirchhoff stress tensors, **P** and **S**, which are derived from a strain energy function as 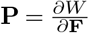 and 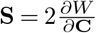, respectively. In the constitutive model presented by Röhrle et al. ^18^, the second Piola–Kirchhoff stress tensor is given by the sum of isotropic, anisotropic and active contributions:

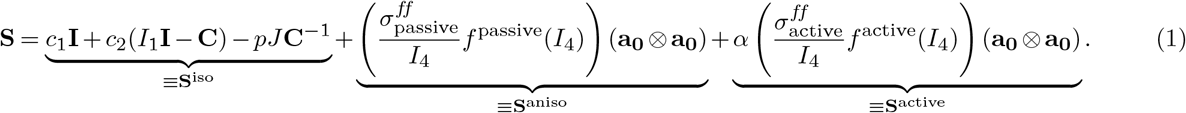

In (1), the isotropic term **S**^iso^ defines a Mooney–Rivlin material with material constants *c*_1_ and *c*_2_ (Table 1) and hydrostatic pressure *p*. The term **S**^aniso^ accounts for passive fiber anisotropy, while **S**^active^ represents active fiber contraction.

**Table 1.**
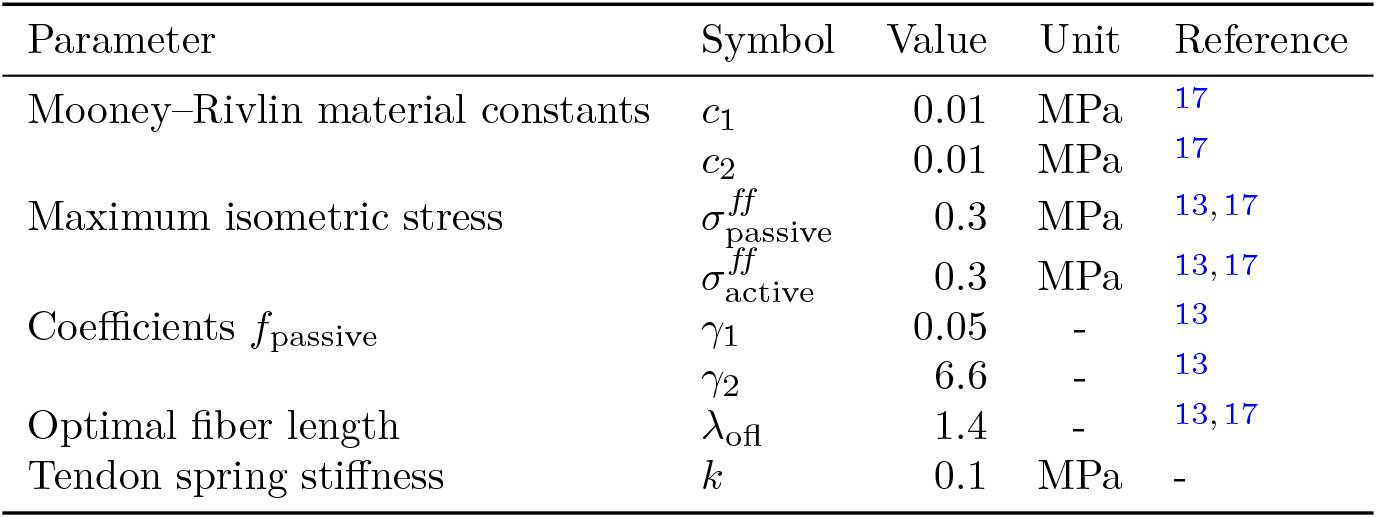
Material constants and force-length relationship parameters for the mechanical muscle tissue model.

The force-length relationships, *f*_passive_ and *f*_active_, depend on the fiber stretch *λ* and follow a standard normalized model^43^ where the maximal active force is generated at an optimal fiber length, *λ*_ofl_ = 1.4 (Figure 2A). Specifically, we define the passive and active components by the following piecewise exponential and polynomial functions, respectively^13^:

**Figure 2.**
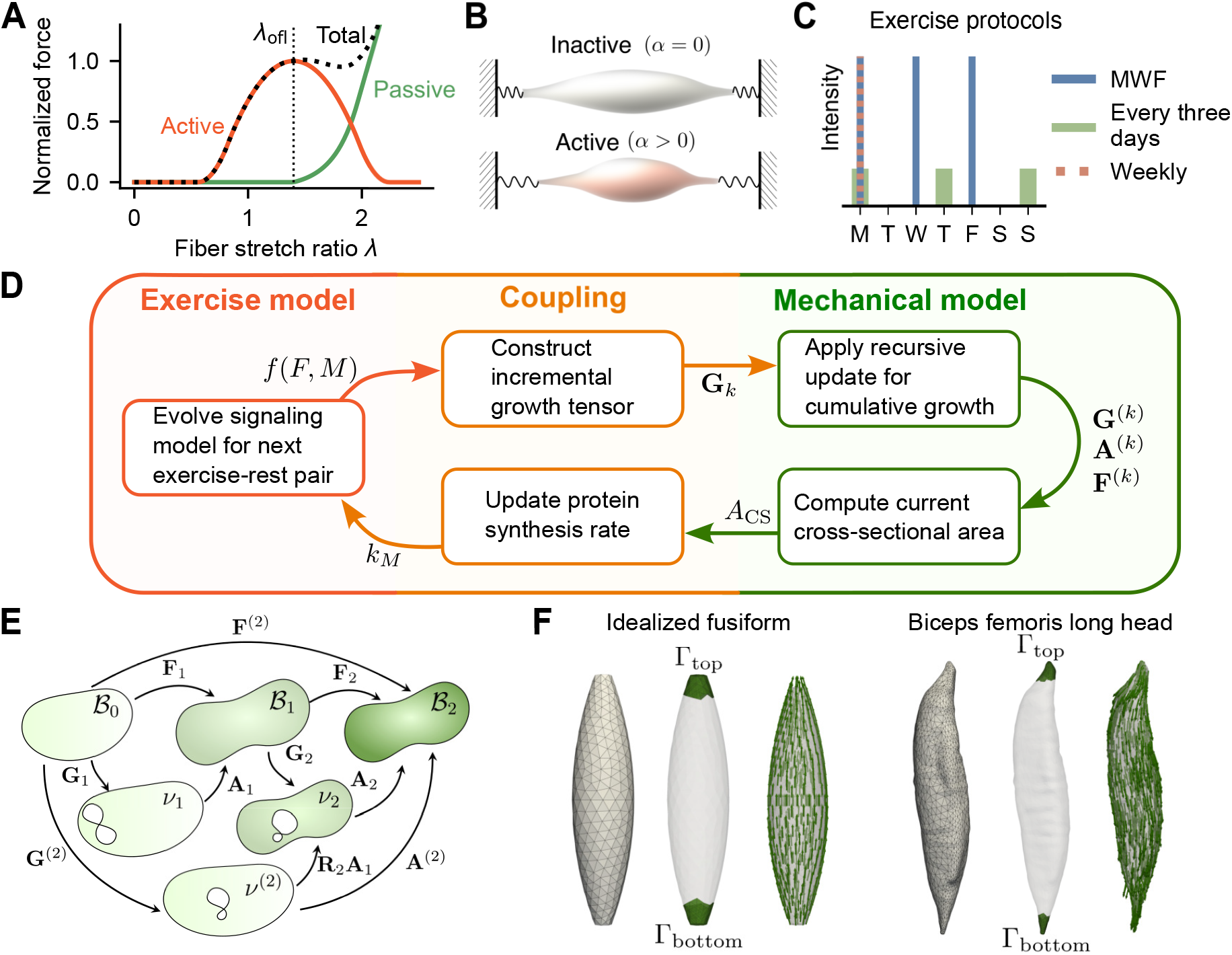
Computational model components and workflow. (**A**) The passive and active fiber force are dependent on stretch *λ*. (**B**) A muscle contraction, showing the inactive (*α* = 0) and active (*α >* 0) states. (**C**) Illustration of one week of the three exercise protocols used in growth simulations. (**D**) The coupling scheme between signaling and growth mechanics used in our computational framework. The exercise model determines an incremental growth tensor **G**_*k*_, which updates the mechanical model through the cumulative growth framework. The resulting cross-sectional area *A*_*CS*_ feeds back to modulate the signaling model via the protein synthesis rate *k*_*M*_ . (**E**) Schematic of the cumulative growth framework, based on the multiplicative decomposition of deformation, **F** = **AG**, and unloading between growth steps. (**F**) Example geometries showing the computational mesh, surfaces for boundary conditions (Γ_top_, Γ_bottom_), and prescribed fiber architecture.

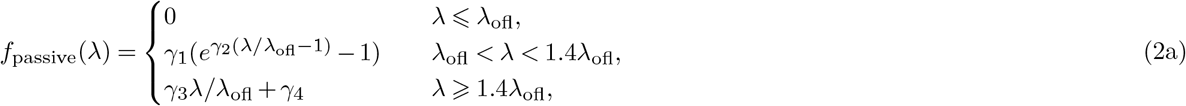

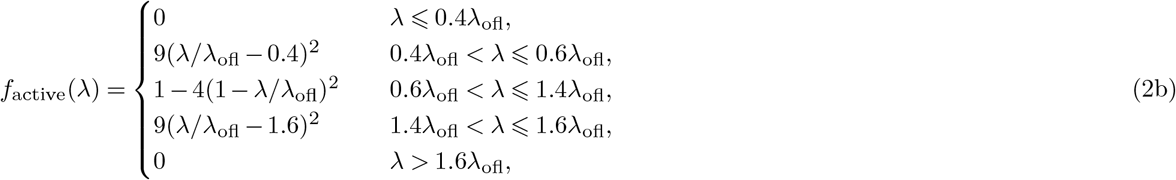

where the parameters *γ*_1_ and *γ*_2_ are given in Table 1, while *γ*_3_ and *γ*_4_ are chosen to ensure continuity and differentiability. Upon activation (*α >* 0), the active stress component generates contraction along the fiber direction **a**_0_ (Figure 2B, F).

### Mechanical equilibrium equations

To determine the muscle tissue’s mechanical state for a given activation level *α*, we solve for the displacement-pressure pair (**u**, *p*) satisfying the governing equations of static equilibrium defined relative to the reference domain Ω:

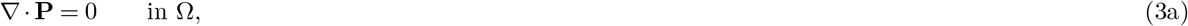

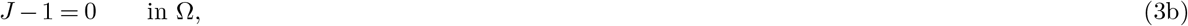

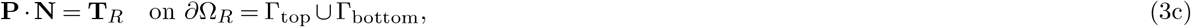

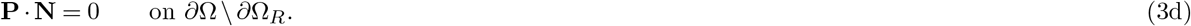

Here, (3a) represents mechanical equilibrium in the absence of body forces, (3b) enforces incompressibility, and **N** is the outward unit normal to the boundary *∂*Ω. The nominal traction **T**_*R*_, entering in the Robin boundary condition (3c), is given as follows. We assume that the physical Cauchy traction **t**, which is defined relative to the deformed boundary surface with unit normal **n**, is negatively proportional to the displacement in the normal direction with proportionality constant *k* ⩾ 0:

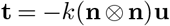

This condition models the tendon(s) connecting the muscle as an elastic spring with stiffness *k* (Figure 2B). We then define **T**_*R*_ as the pull-back of **t** to the reference configuration:

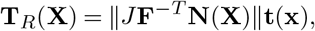

where we have used Nanson’s formula to map between area in the reference and deformed configurations. The remaining boundaries are traction-free (3d).

### Signaling model

The IGF1-AKT-mTOR-FOXO signaling pathway is a central regulator of muscle mass, balancing protein synthesis and degradation^7^. To model the biochemical drivers of hypertrophy, we adopt the signaling model from Villota-Narvaez et al^44^, which in turn builds upon a simplified representation of the pathway by Schiaffino and Mammucari^21^ (Figure 1B). The model consists of a system of ordinary differential equations (ODEs) describing the pathway’s key components. Let *I, A, F*, and *M* denote the normalized populations of IGF1, AKT, FOXO, and mTOR, respectively. Their dynamics are modeled using a Lotka–Volterra framework that captures the activating and inhibiting interactions between them through intrinsic growth rates (*a*_*i*_), self-inhibition rates (*b*_*i*_), and coupling strengths (*c*_*ij*_) (Table 2):

**Table 2.** Baseline parameter values for the IGF1-AKT-mTOR-FOXO signaling model, adapted from^44^, with 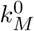 calibrated, and initial conditions adjusted to equilibrium values.

| Parameter | Symbol | Value | Unit |
| --- | --- | --- | --- |
| Intrinsic growth rate | $a_I^0$ | $9.000 \times 10^{-2}$ | $\text{h}^{-1}$ |
| | $a_A^0$ | $4.875 \times 10^{-1}$ | $\text{h}^{-1}$ |
| | $a_F$ | $1.068 \times 10^{-2}$ | $\text{h}^{-1}$ |
| | $a_M$ | $4.635 \times 10^{-3}$ | $\text{h}^{-1}$ |
| Self-inhibition rate | $b_I$ | 2.0 | $\text{h}^{-1}$ |
| | $b_A$ | $5.000 \times 10^{-1}$ | $\text{h}^{-1}$ |
| | $b_F$ | $2.000 \times 10^{-2}$ | $\text{h}^{-1}$ |
| | $b_M$ | $1.000 \times 10^{-2}$ | $\text{h}^{-1}$ |
| Coupling strength | $c_{AI}$ | $2.846 \times 10^{-1}$ | $\text{h}^{-1}$ |
| | $c_{FA}$ | $2.000 \times 10^{-3}$ | $\text{h}^{-1}$ |
| | $c_{MA}$ | $1.139 \times 10^{-3}$ | $\text{h}^{-1}$ |
| | $c_{MF}$ | $2.500 \times 10^{-3}$ | $\text{h}^{-1}$ |
| Min. myofibril population | $N_{\min}$ | 0.5 | - |
| Max. myofibril population | $N_{\max}$ | 1.3 | - |
| Threshold value FOXO | $F^*$ | 0.434 | - |
| Threshold value mTOR | $M^*$ | 0.469 | - |
| Protein synthesis rate | $k_M^0$ | $1.000 \times 10^{-2}$ | $\text{h}^{-1}$ |
| Protein degradation rate | $k_F$ | $1.900 \times 10^{-2}$ | $\text{h}^{-1}$ |
| $a_I(t)$ scale parameter | $\eta$ | 0.04 | $(\text{h} \cdot \text{kgf})^{-1}$ |
| $a_I(t)$ max. muscle force | $F_{\max}$ | 150 | kgf |
| $a_A(t)$ scale parameter | $\tau_s$ | 6 | h |
| $a_A(t)$ delay parameter | $\tau_d$ | 17 | h |
| Initial condition IGF1 | $I_0$ | $4.500 \times 10^{-2}$ | - |
| Initial condition AKT | $A_0$ | 1.001 | - |
| Initial condition FOXO | $F_0$ | $4.339 \times 10^{-1}$ | - |
| Initial condition mTOR | $M_0$ | $4.690 \times 10^{-1}$ | - |
| Initial condition myofibrils | $N_0$ | 1.001 | - |

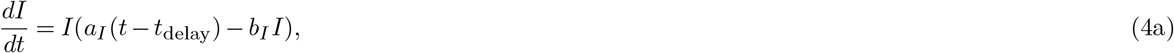

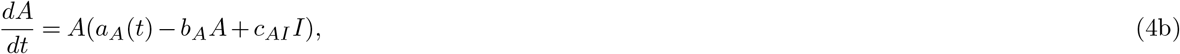

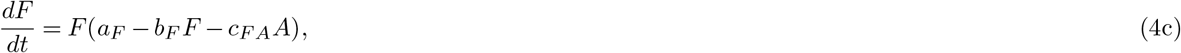

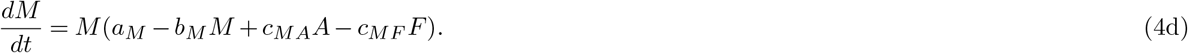

The rate of change of the myofibril population *N* reflects a balance between the mTOR-driven synthesis and FOXO-driven degradation:

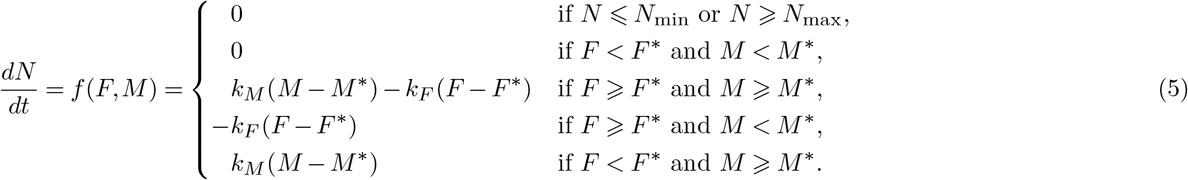

This balance is driven by the protein synthesis rate *k*_*M*_ (Supplementary Figure 2) and degradation rate *k*_*F*_, with contributions effective only when mTOR and FOXO exceed their respective thresholds, *M* ^*∗*^ and *F* ^*∗*^. The myofibril population is also constrained to lie between *N*_min_ and *N*_max_.

Exercise is modeled as a stimulus that modulates the intrinsic growth rates of both IGF1 and AKT. First, the IGF1 growth rate, *a*_*I*_ (*t*), is elevated during exercise according to the relation

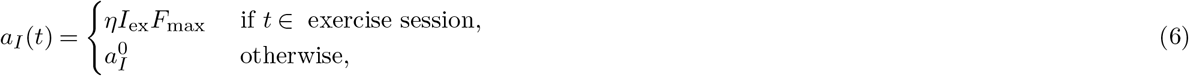

where *η* is a scaling factor, *I*_ex_ ∈ [0, 1] is the exercise intensity factor, *F*_max_ is the muscle’s maximum force capacity, and 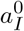 denotes the baseline rate (Table 2). The stimulus is subject to a physiological delay, which we represent by *t*_delay_ in the governing equation for IGF1 (4a) and set to *t*_delay_ = 12h. Second, the intrinsic growth rate of AKT, *a*_*A*_(*t*), responds to exercise with a transient increase defined by

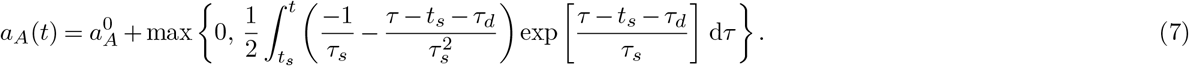

In this expression, *τ*_*d*_ is a delay parameter, *τ*_*s*_ a time-scaling factor, and *t*_*s*_ marks the start time of the most recent exercise session. This formulation creates a transient increase in AKT’s growth rate that decays over several hours, while the max operation ensures *a*_*A*_(*t*) does not fall below its baseline level, 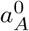.

#### Exercise protocols

We simulated three distinct exercise regimens to test protocol-dependent adaptation (Figure 2C). These were: a high-intensity, high-frequency protocol (1h at 80% intensity, 3x/week, MWF) based on DeFreitas et al.^45^; a high-intensity, low-frequency protocol (1h at 80% intensity, 1x/week); and a low-intensity, medium-frequency protocol (4h at 20% intensity, every three days). The intensity factor *I*_ex_ was set to 0.8 and 0.2 for the high-intensity and low-intensity regimes, respectively. For all protocols, there is an initial week without exercise to establish baseline conditions, followed by eight weeks of the specified exercise regimen.

#### Initial conditions

The initial conditions (*I*_0_, *A*_0_, *F*_0_, *M*_0_, *N*_0_) for the signaling model represent a homeostatic, pre-exercise steady state. To determine these values, we simulated the ODE system (4) with baseline parameter values and no external stimulus 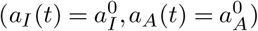 until a stable equilibrium was reached. These equilibrium values, listed in Table 2, were used for all subsequent simulations.

#### Spatial variations in signaling

Spatial heterogeneity and inter-individual variability are emulated by perturbing selected signaling parameters locally. Each baseline parameter *p*_0_ is replaced by a spatially varying parameter

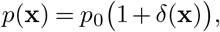

where *δ*(**x**) is a random field sampled from a uniform distribution on [−Δ, Δ] (e.g. Δ = 0.1 for ±10% variation). Perturbations were drawn independently per mesh cell to represent within-muscle heterogeneity, and sampled once per simulation to simulate between-subject variability.

#### Sensitivity analysis

We performed a Sobol sensitivity analysis^46^ using the SALib Python library^47, 48^ to identify the most influential parameters in the signaling model. The final myofibril population (*N* ) after a 15-day simulated exercise protocol was selected as the primary output metric, serving as a computationally efficient proxy for muscle growth. For the analysis, each parameter was sampled from a uniform distribution spanning 50% to 200% of its baseline value. To ensure a robust assessment, the analysis included first- and second-order effects with 10,000 samples, resulting in 300,000 total model evaluations.

### Model coupling

To capture the interplay between molecular signaling and tissue mechanics during skeletal muscle hypertrophy, we couple the subcellular signaling pathway with the macroscopic mechanical model. This integrated framework enables simulation of long-term muscle adaptation, where molecular events and mechanical states dynamically influence each other. We implement this bidirectional coupling through two primary mechanisms: (1) the signaling dynamics inform a volumetric growth tensor that drives changes in muscle geometry, and (2) the resulting change in muscle size provides feedback to the signaling pathway by modulating a key parameter – the protein synthesis rate (Figure 2D). These two coupling mechanisms are detailed in the following sections.

#### From signaling to macroscopic growth through the growth tensor

Molecular signals are translated into macroscopic growth by linking the myofibril population *N* (*t*) from the signaling model to the tissue’s deformation. This coupling is formalized using the theory of volumetric growth, where the total deformation gradient **F** is decomposed into a growth part **G** and an elastic part **A**, such that **F** = **AG**^29, 31^. In this decomposition, the growth tensor **G** represents the local (potentially incompatible) addition of mass driven by biology, while **A** is the subsequent elastic deformation required to maintain mechanical equilibrium and ensure the tissue remains a compatible, continuous body.

To determine **G**, we first calculate a scalar growth multiplier, *G*(*t*). The core assumption linking *N* (*t*) to the continuum growth is that the myofibril population is directly proportional to the muscle cross-sectional area (CSA), *A*_*CS*_(*t*), via a constant *κ*, such that *N* (*t*) = *κA*_*CS*_(*t*). From the change in myofibrils Δ*N* = *f* (*F, M* )Δ*t* predicted by the signaling model over a time step Δ*t*, the growth multiplier is defined as:

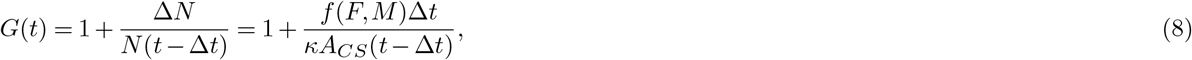

where *N* (*t −* Δ*t*) is the myofibril population at the previous time step. This scalar multiplier then defines the anisotropic growth tensor **G**:

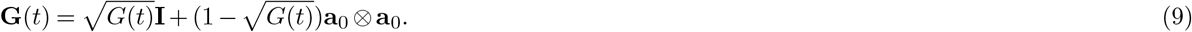

This formulation ensures that growth occurs exclusively in the plane perpendicular to the fiber direction **a**_0_, consistent with an increase in CSA^32^.

#### Feedback from mechanics to signaling

The bidirectional coupling is completed by a feedback mechanism where the mechanical state of the muscle modulates the signaling pathway. Concretely, the protein synthesis rate *k*_*M*_ is set to depend on the current CSA of the muscle *A*_*CS*_. We define the normalized CSA as 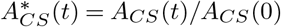, where *A*_*CS*_(0) is the initial area. The synthesis rate is then updated as follows:

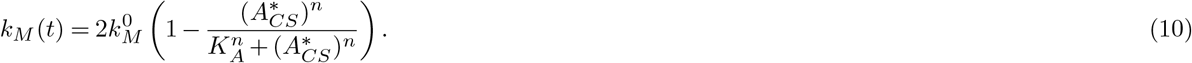

With a Hill coefficient of *n* = 4 and a half-max constant of *K*_*A*_ = 1, this function establishes homeostatic control. The synthesis rate returns to its baseline value when 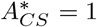, decreases during hypertrophy 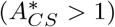 to limit excessive growth, and increases during atrophy 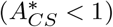 to stimulate growth back towards the baseline. This bounded, Hill-type regulation provides a simple, yet physiologically plausible feedback loop for muscle size adaptation (Supplementary Figure 3).

#### Implementation of cumulative growth

To model the long-term effects of incrementally applied growth, we adopt the cumulative growth framework of Goriely and Ben Amar^38^. A key requirement of this framework is that each new growth increment, **G**_*k*_, must be applied to a stress-free configuration. Because the tissue is generally under residual stress from previous growth steps, this requires computationally “unloading” the accumulated elastic deformation before applying the next increment (Figure 2E). After *k* growth steps, the total deformation **F**^(*k*)^ is decomposed into a total elastic tensor **A**^(*k*)^ and a total growth tensor **G**^(*k*)^, such that **F**^(*k*)^ = **A**^(*k*)^**G**^(*k*)^. These total tensors are updated recursively at each step *k*:

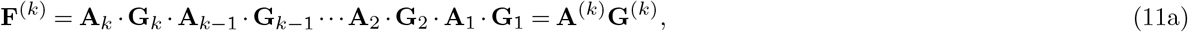

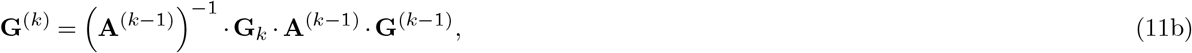

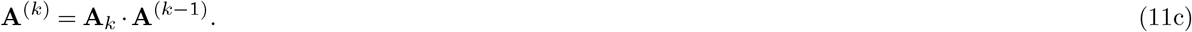

In this formulation, the incremental growth tensor **G**_*k*_ is determined by the signaling model via (9). Equation (11b) integrates this new growth by first computationally unloading the previous state via (**A**^(*k−*1)^)^*−*1^. The new incremental elastic tensor **A**_*k*_ is found by solving for mechanical equilibrium after the growth has been applied, and the total elastic deformation is then updated via (11c).

### Muscle geometries

#### Mesh generation

Simulations were performed on three distinct types of computational geometries: a cylindrical geometry, an idealized fusiform muscle, and a set of anatomically realistic muscle models (Figure 2F). The idealized fusiform geometry was created to represent a simplified, yet physiologically relevant, muscle shape. The tetrahedral mesh was generated using Gmsh^49^ and consists of 1997 cells. The realistic muscle geometries were derived from segmentations of the Visible Human Dataset^39, 40^. From these surface representations, tetrahedral meshes were generated using fTetWild^50^, resulting in cell counts between 6707 and 9668.

#### Muscle fiber architecture

Defining the muscle fiber architecture, which sets the local anisotropy **a**_0_, is essential for all geometry types. For the cylinder, fibers simply run parallel along the long axis. In the idealized fusiform model, fibers were defined analytically via a spline curve. For the realistic geometries, fiber directions were estimated by solving a Stokes flow problem^51^. This involved setting the muscle origin and insertion as inflow and outflow boundaries, applying slip conditions on the remaining surface, and normalizing the resulting velocity field to define the local fiber vectors (Figure 2F).

#### Muscle cross-sectional area

The cross-sectional area (CSA) was computed by integrating over a transverse reference surface Γ_*CS*_ intersecting the muscle at the midpoint of its longitudinal axis. Nanson’s formula^52^ provides the mapping from the reference to the deformed area element for this integration:

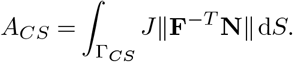

For the idealized geometry, the surface Γ_*CS*_ was a disc embedded directly into the 3D mesh during its generation with Gmsh. For the realistic geometries, the orientation of this surface was determined using principal component analysis (PCA), defining Γ_*CS*_ as the plane passing through the geometric centroid of the mesh and perpendicular to the muscle’s primary longitudinal axis. The displacement field was interpolated onto this plane to compute the deformed area.

### Numerical solution schemes and verification

To simulate the coupled mechanics-signaling system, we use a quasi-static approach, justified by the slow timescale of hypertrophy compared to that of elastic deformation^38^. The simulation advances through a series of exercise and rest periods, using a time step of Δ*t* = 1 h. Each period begins by solving the signaling ODE system to determine the rate of change in the myofibril population. This growth is then applied incrementally in a nested loop, where each growth step consists of constructing a new growth tensor, solving for mechanical equilibrium, and finally, closing the feedback loop by updating the protein synthesis rate based on the current cross-sectional area (Supplementary Algorithm 1). The computational model was implemented in Python, and the simulation code is openly available^53^.

The signaling ODE system was integrated using the ‘RK45’ method from SciPy’s `solve_ivp` function^54^, with a local time step of dt = 0.05h. For the spatially varying signaling models, however, a custom implementation of the fourth-order Runge–Kutta (RK4) method was used. We use the finite element method to numerically solve the nonlinear mechanical equilibrium equations (3), using the FEniCSx finite element software^55^. Relative to each of the computational meshes, we define the finite element space **V** as the space of continuous piecewise quadratic 3-vector fields and *Q* as the space of continuous piecewise linears. At each iteration, we solve the following system of nonlinear equations: find the approximate displacement **u** *∈* **V** and pressure *p ∈ Q* solving

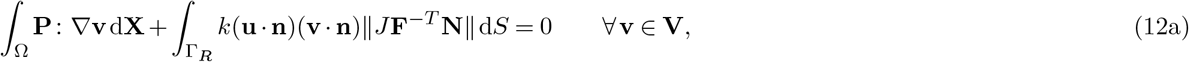

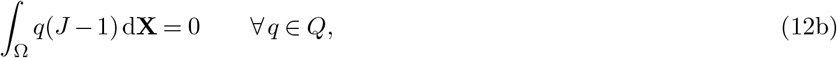

where **P** is the Piola–Kirchhoff tensor cf. (1). The nonlinear system (12) was solved using a Newton solver. To verify the implementation, we performed a series of numerical tests, including evaluating the correctness of the numerical solution algorithms for the spatially varying ODE systems (Supplementary Figure 1).

## Results

### Signaling model links exercise protocol to distinct dynamic responses

Before implementing the full multiscale framework, we first characterized the dynamics of the core signaling model in isolation. We simulated the model’s response to three distinct exercise protocols (Figure 2C): (1) a high-intensity, high-frequency protocol (MWF); (2) a high-intensity, low-frequency protocol (weekly); and (3) a low-intensity, medium-frequency protocol (every three days).

The model exhibited a dynamically consistent response across all protocols (Figure 3A-E). Each exercise bout triggered a rapid, transient spike in the upstream signals IGF1 and AKT, with a peak observed approximately 12 hours post-stimulus (Figure 3A-B). The amplitude of these spikes was directly proportional to the exercise intensity. The upstream activation drove the expected opposing response in the downstream effectors: mTOR levels (promoting synthesis) increased, while FOXO levels (promoting degradation) were suppressed (Figure 3C-D). Importantly, the system operates on two distinct timescales. IGF1 and AKT act as fast responders, returning to baseline between each exercise period. In contrast, FOXO and mTOR display a wave summation effect, wherein the responses to repeated stimuli accumulate and oscillate around an apparent steady state. The magnitude of this shift is protocol-dependent and directly determines the final myofibril density *N* (Figure 3E).

**Figure 3.**
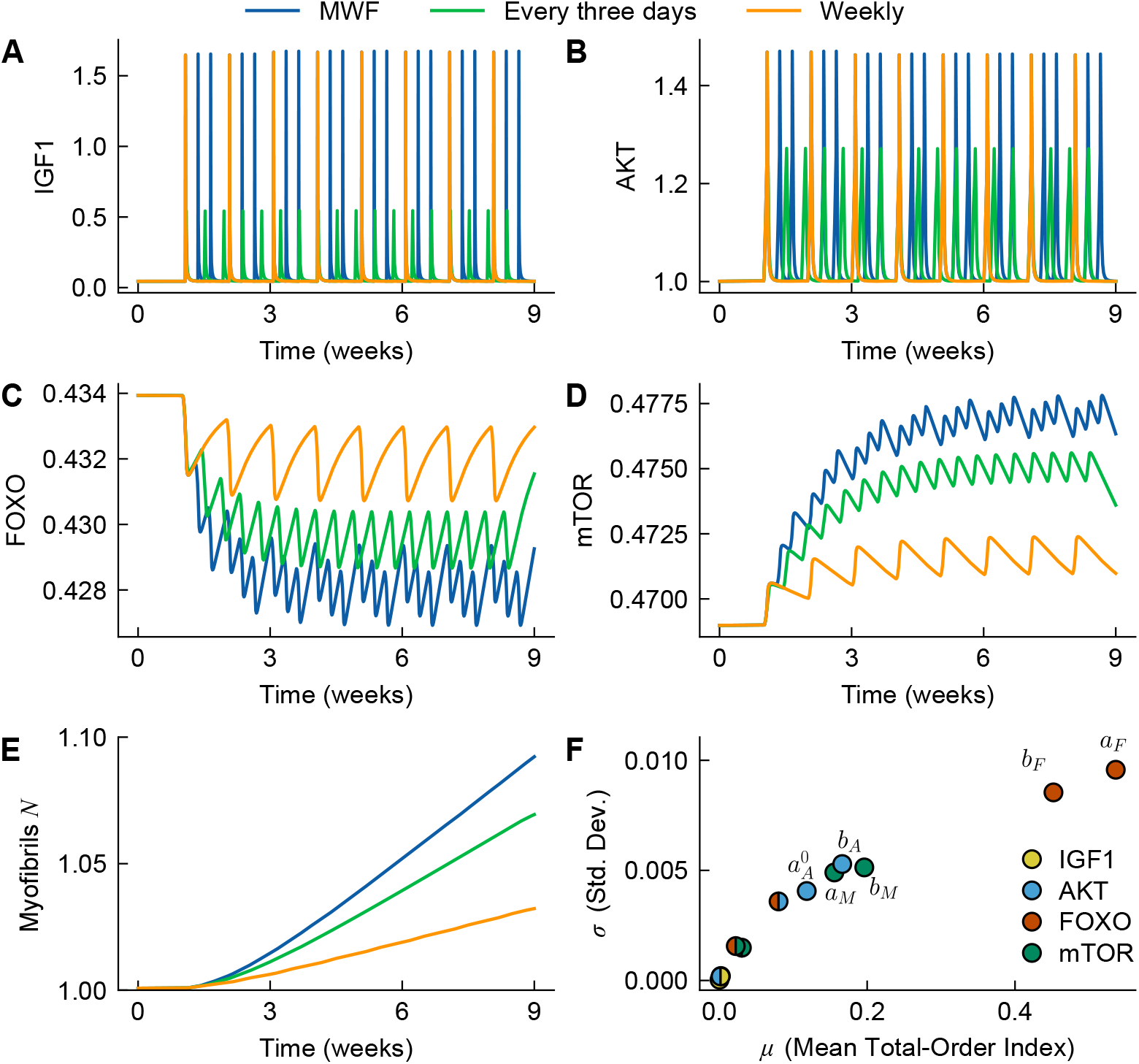
Dynamics of the IGF1-AKT signaling model in response to different exercise protocols. (**A–E**) Time courses of (**A**) IGF1, (**B**) AKT, (**C**) FOXO, (**D**) mTOR, and (**E**) myofibrils *N* for three different exercise protocols. (**F**) Sobol sensitivity analysis quantifying the influence of each model parameter on the final myofibril number obtained using a standardized daily exercise protocol. The plot shows the mean Sobol index versus its standard deviation for each parameter. Marker colors identify the associated signaling molecule, with split colors indicating coupling parameters.

To identify which model parameters were most critical in controlling this response, we performed a Sobol sensitivity analysis. The analysis revealed that the final myofibril population was most sensitive to parameters governing the downstream effector, FOXO (Figure 3F). Specifically, the intrinsic growth (*a*_*F*_ ) and self-inhibition (*b*_*F*_ ) rates of FOXO were the most influential, followed by the corresponding rates for mTOR (*a*_*M*_ and *b*_*M*_ ) and AKT (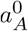 and *b*_*A*_). This sensitivity arises because the core signaling network functions as an incoherent feedforward loop^23, 24^: the upstream signal (AKT) simultaneously promotes a synthesis pathway (mTOR) and suppresses a degradation pathway (FOXO). Consequently, the qualitative outcome – whether the muscle undergoes hypertrophy or atrophy – is not determined by the exercise protocol alone but depends critically on the kinetic balance between these opposing branches. The high sensitivity to FOXO parameters indicates that *a*_*F*_ and *b*_*F*_ effectively set a threshold for growth. Small perturbations to these rates can abruptly toggle the net protein turnover from positive to negative, even under identical exercise stimuli.

Comparing the net growth response between the regimens, the high-intensity, high-frequency (MWF) protocol induced the largest increase in the myofibril population (*N*) due to the summation effects described above (Figure 3E). However, all exercise protocols predicted net growth relative to the sedentary baseline, confirming that even lower-frequency stimuli provide a sufficient perturbation to shift the net protein balance. With these protocol-dependent trajectories of *N* established, we next used them as the driving input for the full 3D framework to investigate how biochemically driven growth manifests as mechanical remodeling across idealized muscle geometries.

### Coupled model predicts protocol-dependent hypertrophy in an idealized muscle geometry

While biochemical signaling and tissue mechanics are often modeled in isolation, physiological growth is an inherently coupled phenomenon. Accordingly, we next integrated the signaling model with the 3D hyperelastic framework, initially simulating long-term adaptation in an idealized fusiform muscle geometry (Figure 4A).

**Figure 4.**
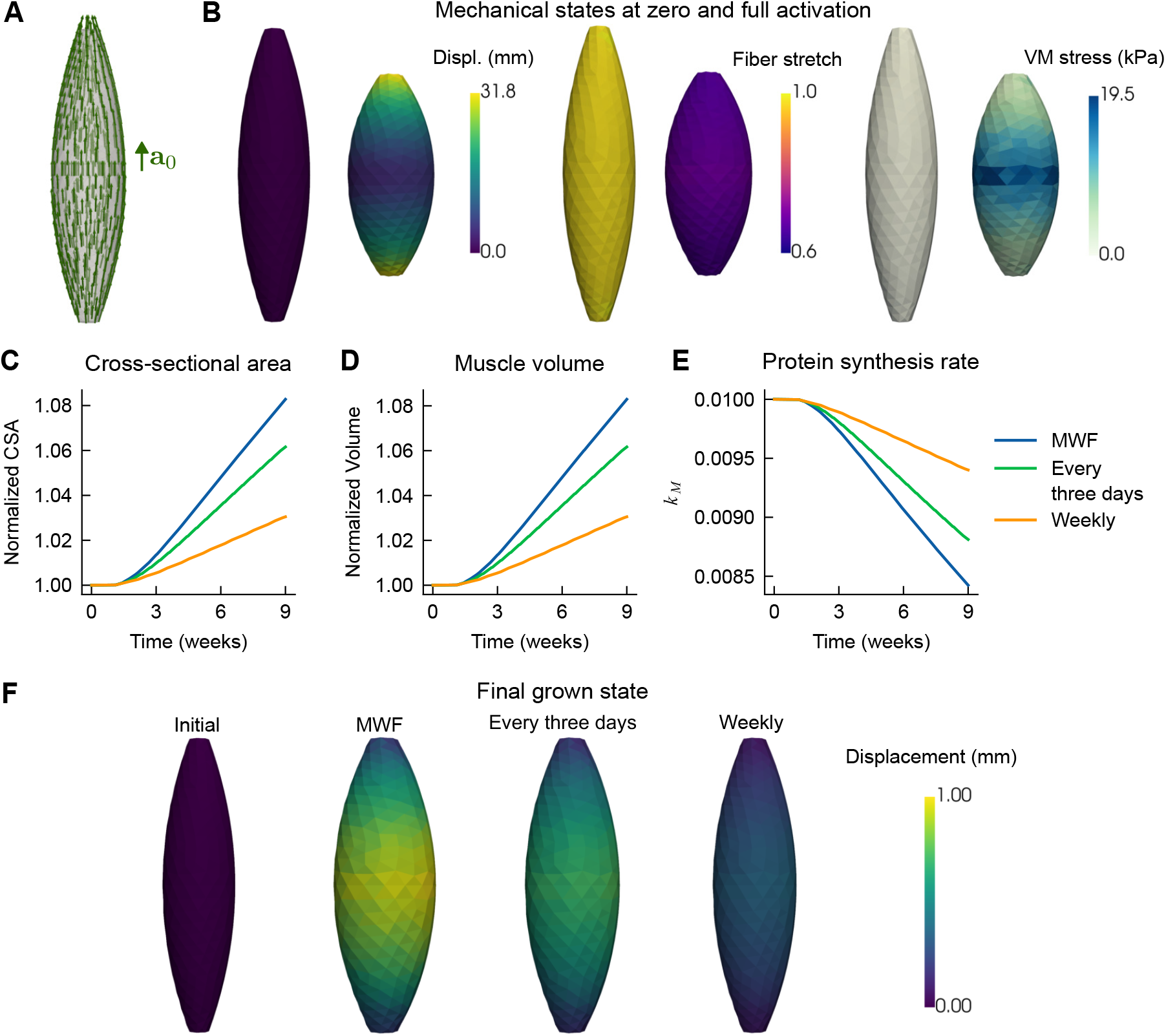
Coupled model simulation of exercise-induced hypertrophy in an idealized fusiform muscle. (**A**) Fiber orientation *a*_0_ computed from a spline expression. (**B**) Mechanical state variables (displacement, fiber stretch *λ*, and von Mises stress) at zero and full activation (*α* = 1). (**C–E**) Predicted time courses of cross-sectional area, muscle volume, and the protein synthesis rate *k*_*M*_ in response to three distinct exercise protocols. (**F**) Final muscle configurations after the simulation period for each protocol, with the growth-induced displacement amplified 10x for visualization.

First, we characterized the muscle’s baseline mechanical behavior in the absence of volumetric growth. A simulated contraction from zero to full activation (*α* from 0 to 1) produced heterogeneous distributions of displacement, stress and fiber stretch along the muscle geometry (Figure 4B). Muscle ends were allowed to partially contract towards the midplane, mimicking the compliance of tendons. The displacement magnitudes were largest at the ends of the muscle, consistent with this shortening behavior. The von Mises stress was concentrated at the center of the muscle, whereas the fiber stretch *λ* was relatively uniformly distributed, exhibiting only slightly higher values in the central region.

Having established that the mechanical framework gives the expected response for an idealized muscle geometry, we next combined the signaling model with the muscle growth model. We simulated muscle adaptation over a nine-week period for the three different exercise protocols. The coupled model successfully translated the protocol-dependent signaling dynamics into different macroscopic growth in an exercise-dependent manner. Consistent with the myofibril-level predictions, the high-intensity, high-frequency MWF protocol induced the most significant increase in both cross-sectional area (CSA) and total muscle volume, followed by the every-three-days and weekly protocols (Figure 4C-E). Furthermore, we also identified the critical role of homeostatic control generated by the feedback between signaling and mechanics. As the muscle CSA increased in response to training, the mechanical feedback loop progressively downregulated the protein synthesis rate, *k*_*M*_, from its baseline value (Figure 4C,E). The strength of this feedback is dependent on the exercise modality – higher exercise frequency leads to a sharper drop in the protein synthesis rate, naturally limiting the rate of further hypertrophy (Figure 4D). The macroscopic effect of this adaptation is visualized in Figure 4F, which shows the final deformed muscle shapes for each protocol (growth amplified tenfold for visualization). This result is a direct consequence of the anisotropic growth tensor included in the model (9), which mechanistically links the signaling output to mass addition in the plane perpendicular to the fiber axis. Overall, these simulations suggest that while any exercise stimulus promotes growth relative to the baseline, the mechanical feedback loop prevents a linear dose-response, ensuring the higher-frequency training leads to self-limiting adaptation rather than unbounded hypertrophy.

### Anatomically realistic models show protocol-dependent adaptation and non-uniform growth

Thus far, we used idealized geometries to investigate how muscle growth responds to different exercise modalities. To determine how complex, subject-specific anatomy influences this response, we next applied the coupled framework to a cohort of 12 anatomically realistic muscle geometries derived from the Visible Human Dataset^39, 40^. The cohort consisted of bilateral (left and right) meshes for both a male and female subject, yielding four unique geometries for each of the three selected muscle types: the biceps femoris long head (BFLH), semitendinosus (ST), and tibialis anterior (TA) (Figure 5A). Functionally, the BFLH and ST act as hamstring knee flexors and hip extensors, while the TA serves as a primary ankle dorsiflexor. These muscles are heavily recruited during common activities such as running, squatting, and lunging, making them highly relevant for investigating exercise-induced adaptation. Prior to simulation, we characterized the baseline geometries of the cohort, confirming that male muscles displayed larger initial volumes and cross-sectional areas compared to their female counterparts (Figure 5B). Furthermore, to capture the internal architecture essential for anisotropic growth, we generated subject-specific fiber arrangements for each mesh using Stokes flow simulations (see Methods and Figure 5C).

**Figure 5.**
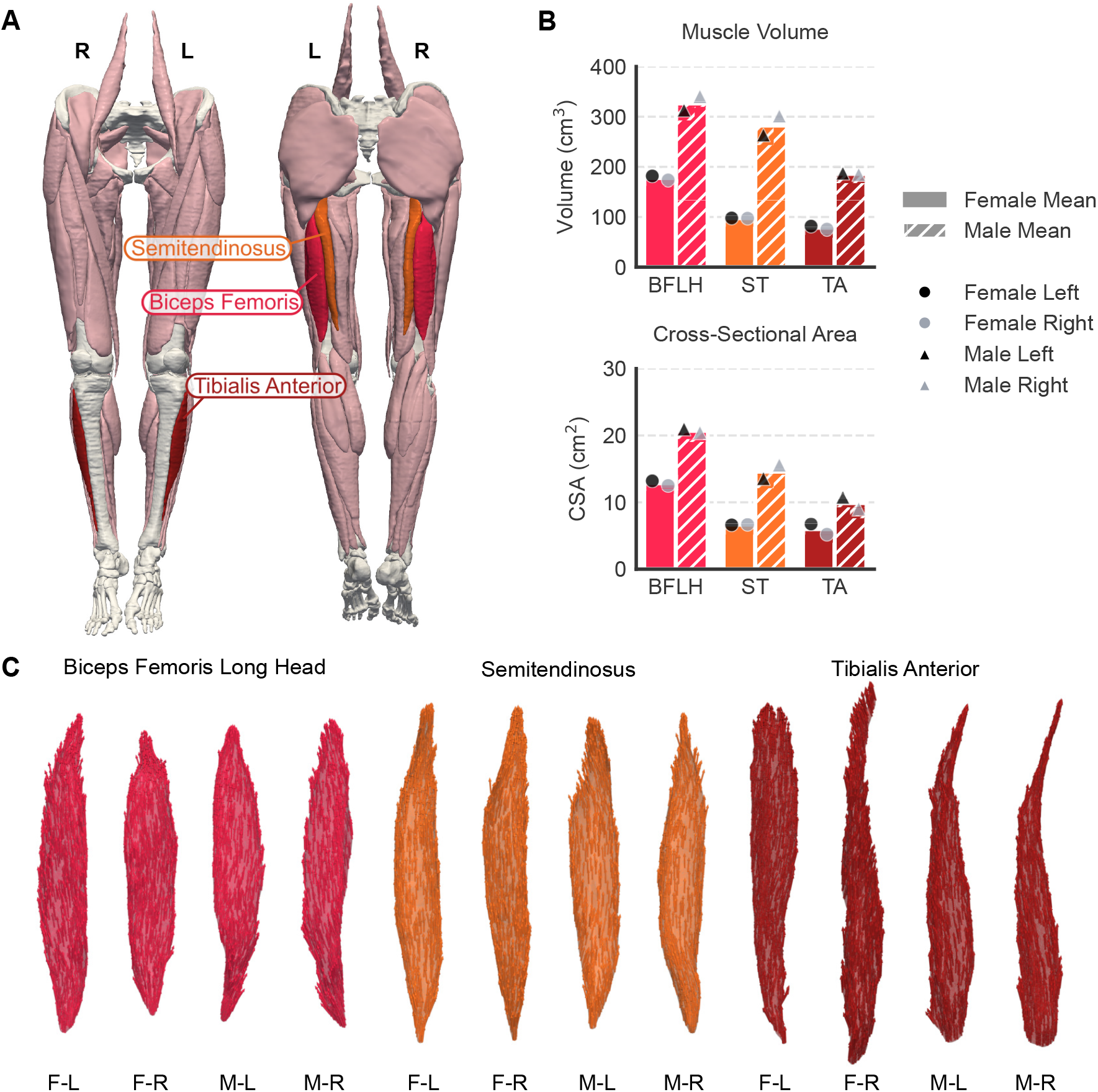
Realistic muscle geometries derived from the Visible Human dataset. (**A**) Anatomical placement of the semitendinosus (ST), biceps femoris long head (BFLH), and tibialis anterior (TA) muscles, visualized within the Visible Human Female. (**B**) Comparison of computed muscle volumes and cross-sectional areas for the twelve meshes derived from the bilateral ST, BFLH, and TA muscles of the Visible Human Female and Male. (**C**) Fiber arrangements generated using Stokes flow simulations within meshes derived from the surface geometries shown in (A). Labels indicate gender (F: Female, M: Male) and laterality (L: Left, R: Right).

We first examined the model’s predictions for a representative case: the female left biceps femoris long head (BFLH). The 3D visualizations of the muscle following the nine-week training protocols reveal a distinct, protocol-dependent hypertrophic response (Figure 6A). Consistent with the high-intensity nature of the stimulus, the MWF protocol produced the most substantial visible thickening of the muscle belly, followed by the every-three-days and weekly protocols. Our simulations show that the complex, subject-specific architecture results in irregular tissue expansion, in contrast to the symmetric radial scaling observed in the idealized model. While the idealized geometry distributes growth evenly around the central axis (Figure 4F), the realistic geometry exhibits spatially heterogeneous hypertrophy, with peak displacement localized to specific regions of the muscle belly. To understand the regulation underlying this growth, we tracked the temporal evolution of the protein synthesis rate, *k*_*M*_ (Figure 6B). The simulations show a progressive downregulation of *k*_*M*_ from its baseline value as the tissue adapts. This decay reflects the homeostatic feedback loop: as the muscle grows, the mechanical stimulus for further synthesis is reduced, naturally limiting the adaptation rate over time.

**Figure 6.**
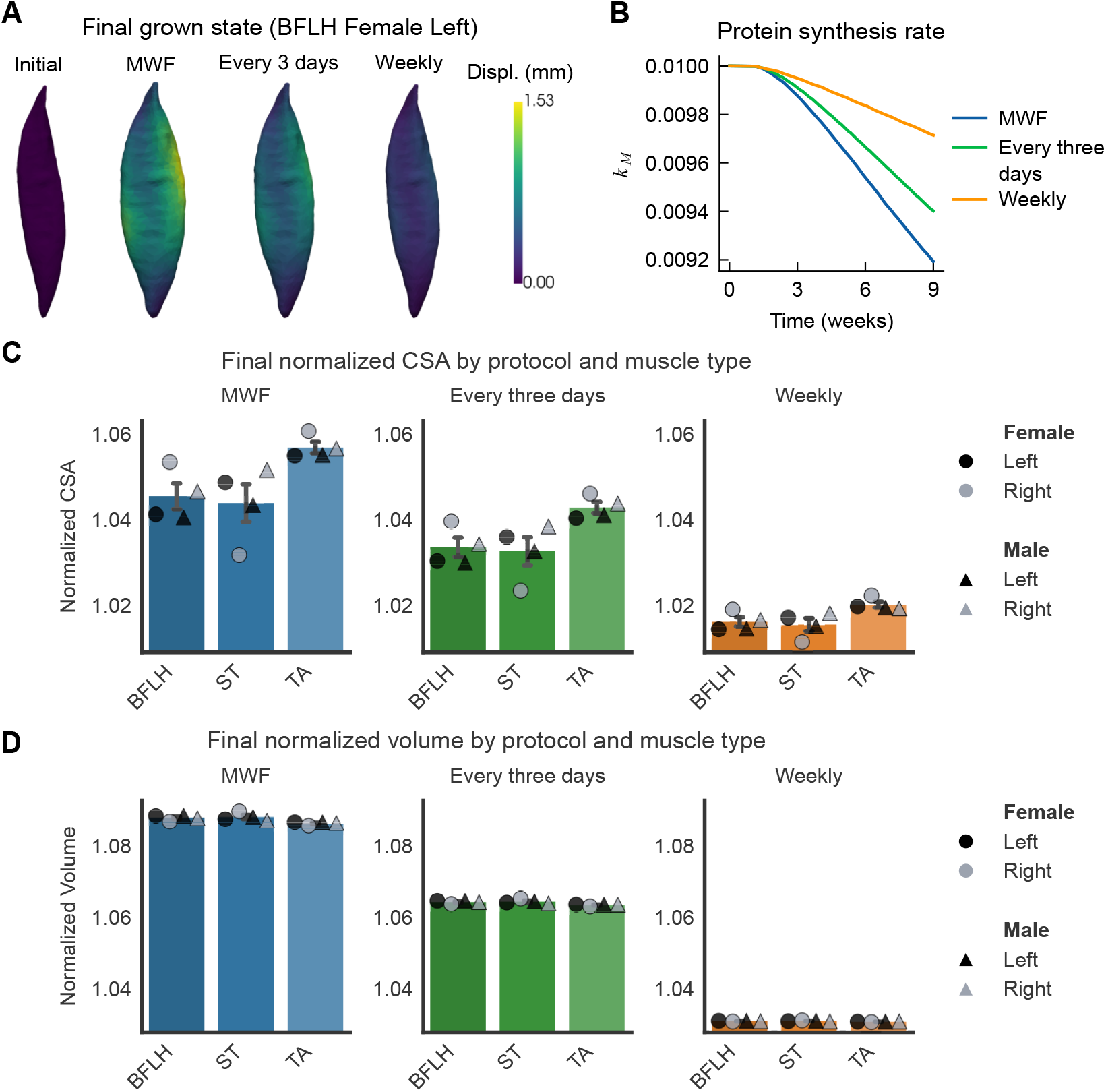
Coupled model simulation of exercise-induced hypertrophy in realistic muscle geometries. (**A**) Final muscle configurations of the female left biceps femoris long head (BFLH) following the simulation period for each exercise protocol. Growth-induced displacements are amplified 10x for visualization. (**B**) Predicted temporal evolution of the protein synthesis rate (*k*_*M*_ ) in response to three distinct loading protocols (shown for the female left BFLH geometry). (**C–D**) Predicted final CSA and volume after three protocols in the BFLH, ST, and TA muscles. Error bars indicate standard error.

Expanding this analysis to the full cohort of 12 geometries, we found that these protocol-dependent trends were robust across all subjects and muscle types. For every geometry tested, the high-frequency, high-intensity MWF protocol induced the most pronounced hypertrophy, followed by the every-three-days and weekly protocols (Figure 6C-D). A key finding from the cohort analysis was that the relative increase in total muscle volume was consistently larger than the relative increase in cross-sectional area (CSA) measured at the muscle’s midplane. For instance, the MWF protocol induced an average volume increase of approximately 8-9% (Figure 6D), while the corresponding CSA increase was only 3-6% (Figure 6C). This discrepancy arises because the complex architecture drives non-uniform growth, where mass is added asymmetrically along the muscle. Consequently, site-specific CSA measurements, a common metric in clinical and experimental studies^56^, may systematically underestimate the extent of tissue adaptation. The variability was notably more pronounced in the normalized CSA measurements (Figure 6C) compared to the more tightly clustered volume measurements (Figure 6D). This reinforces that CSA is a localized metric, highly sensitive to anatomical placement and local geometric irregularities, whereas volume more robustly captures the global adaptation. Within the scope of our simulations, we did not see any muscle type specific or sex specific differences in response to exercise.

In conclusion, these results demonstrate that tissue geometry and fiber architecture play a fundamental role in the spatial distribution of hypertrophy. Because the model defines growth as mass addition transverse to the local fiber axis, the subject-specific fiber orientation directly dictates the direction of expansion at every point. Consequently, complex fiber arrangements result in non-uniform shape changes that cannot be fully captured by idealized models, highlighting the need for anatomical fidelity in predictive modeling of muscle adaptation.

### Tissue mechanics integrates heterogeneous signaling into smooth deformation

Finally, we investigated how inherent spatial variation in biochemical signaling affects the mechanochemical response to exercise. Moving beyond the assumption of homogeneous tissue properties, we conducted an ensemble analysis on the female left BFLH geometry using the high-intensity MWF protocol. This specific combination was selected as the “maximal response” case to determine if robust hypertrophy persists even under noisy signaling conditions. In these simulations, all rate parameters associated with IGF1, AKT, FOXO, and mTOR were simultaneously perturbed by ±1% using a uniform random distribution at each mesh cell. Crucially, given the incoherent feed-forward loop structure of the signaling network, such parameter fluctuations have the potential to qualitatively alter the local outcome, flipping the kinetic balance from net synthesis to degradation.

The time courses of the local ODE states reveal the model’s sensitivity to cellular variability (Figure 7A). While the ensemble’s mean signaling response remained consistent with the uniform baseline, the variance was significantly amplified in the downstream regulators, FOXO and mTOR. This amplification reflects a key biological feature: because downstream molecules integrate signals over time, they accumulate and magnify small upstream fluctuations. Consequently, even under the optimal high-intensity (MWF) stimulus, local outcomes vary significantly. While the average response is positive, specific regions may experience blunted growth or even atrophy due to unfavorable parameter combinations.

**Figure 7.**
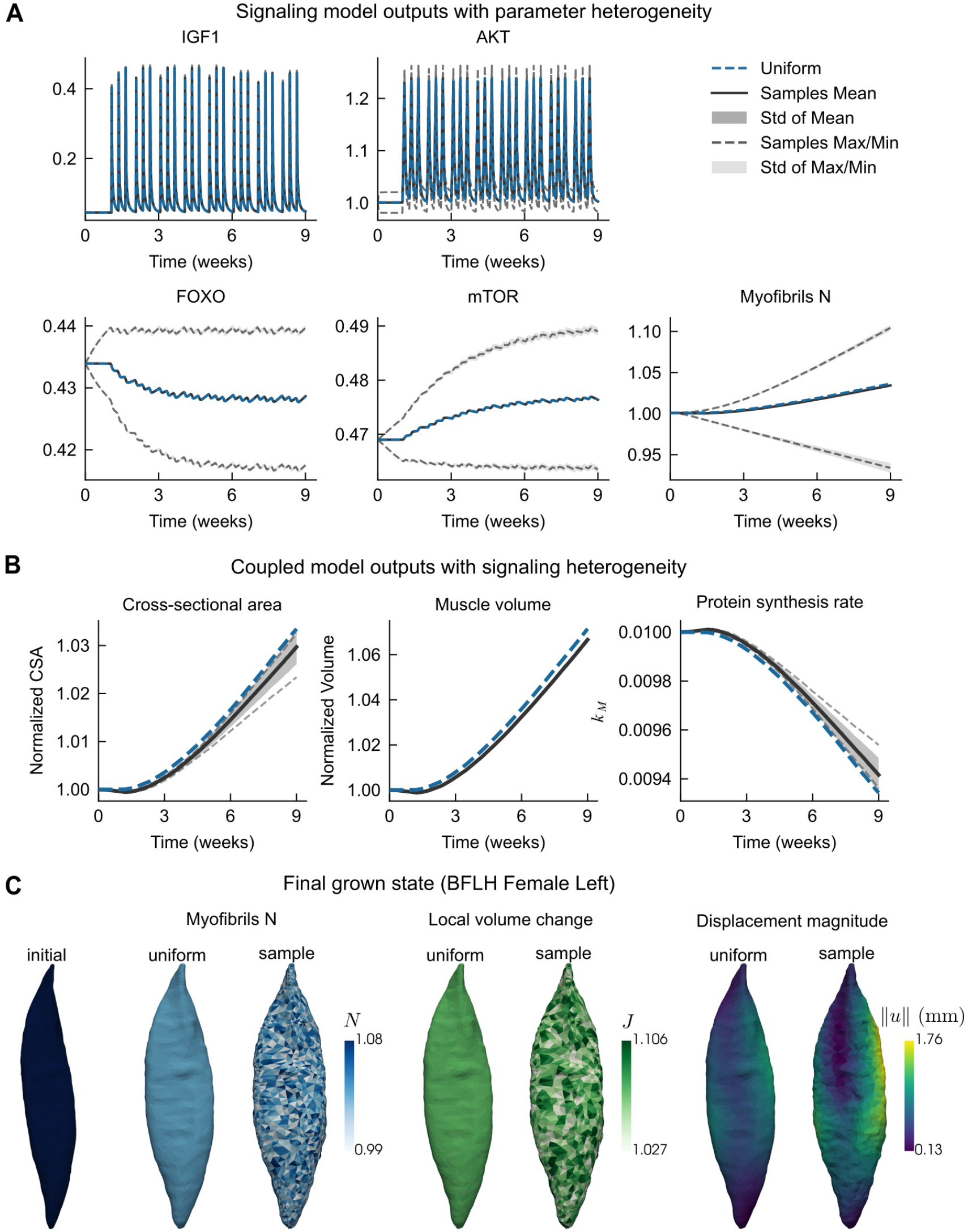
Impact of signaling parameter heterogeneity on local and global growth dynamics. (**A**) Predicted time courses of the local ODE states (IGF1, AKT, FOXO, mTOR, and myofibril population N). Heterogeneity was introduced by sampling rate parameters (*a*_*i*_, *b*_*i*_, *c*_*ij*_) from uniform distributions within ±1% of their baseline values. Gray lines and shading indicate the mean ±1 standard deviation across five samples; the dashed blue line represents the baseline uniform case. (**B**) Predicted evolution of normalized cross-sectional area (CSA), normalized volume, and the protein synthesis rate *k*_*M*_ under the MWF protocol (colors as in **A**). (**C**) Spatial maps of final myofibril density (N), local volume change (*J*), and growth-induced displacement magnitude (amplified 10x) for the uniform case versus a representative heterogeneous sample.

At the macroscopic level, the uniform baseline case consistently achieved more total growth than the mean of the heterogeneous ensemble, as measured by both normalized volume and CSA (Figure 7B). This indicates that the introduction of signaling noise creates a net drag on the system, where the cumulative effect of suboptimal local responses slightly dampens the aggregate adaptation compared to the idealized, uniform condition. This difference in growth is reflected in the protein synthesis rate, *k*_*M*_ . Since the uniform case achieves a larger increase in CSA, it triggers a correspondingly larger decrease in *k*_*M*_ compared to the heterogeneous average. Furthermore, this analysis confirmed our previous observation that variability was consistently greater for normalized CSA than for total volume. Because CSA is a localized slice, it is highly susceptible to specific regional fluctuations, whereas total volume integrates over the entire domain, effectively smoothing out the local noise to provide a more robust metric of adaptation.

The 3D spatial maps reveal a striking contrast between the biological drive and the mechanical response (Figure 7C). The heterogeneity in the myofibril population (*N* ) is closely reflected in the speckled, fragmented map of local growth (*J* ). However, the resulting displacement magnitude shows a more smooth and coordinated pattern. This phenomenon demonstrates a mechanical buffering, where the global equilibrium constraints of the tissue force the continuum to deform smoothly. Effectively, the elastic matrix averages out the highly variable local expansion rates, preventing individual cells from expanding or shrinking in isolation. However, this buffering comes at a cost. The mismatch between the jagged local growth drive (*J* ) and the smooth deformation gradient implies the generation of internal residual stresses. It is possible that regions of muscle tissue attempting to grow rapidly are physically constrained by their slower-growing neighbors, creating a mechanical drag that not only dampens the aggregate expansion but may also generate localized shear concentrations prone to micro-damage.

## Discussion

In this work, we developed a multiscale computational framework to quantify the effects of different exercise modalities on realistic human muscle geometries. Our multiscale computational framework successfully integrated IGF1-AKT signaling dynamics with 3D hyperelastic tissue mechanics to model long-term skeletal muscle hypertrophy. We first established that the readouts of the signaling pathway are highly sensitive to the exercise protocol, with the high-intensity, high-frequency regimen inducing the most significant growth. The architecture of the signaling network resembles an incoherent feedforward loop^23, 24^. We identified FOXO and mTOR as the critical points governing the balance between protein synthesis and degradation for muscle hypertrophy. The same stimulus resulting in a balance of synthesis and degradation is a common feature found in many signaling pathways^57–59^ and is important for maintaining homeostasis. Applying the coupled model to anatomically realistic human geometries confirmed this protocol-dependent adaptation, highlighting that the feedback between mechanical deformation and signaling is essential for regulating the magnitude of hypertrophy and stabilizing the growth process. Our simulations also revealed that the relative increase in total muscle volume consistently exceeds the increase in cross-sectional area (CSA) in realistic geometries. Finally, our analysis of spatial heterogeneity revealed that while local volume expansion rates are highly localized by biological variability, the tissue-wide integration of mechanics acts as a buffer, forcing the overall displacement and resulting bulk shape change to remain relatively smooth and coordinated.

Our computational framework aligns well with established anatomical and physiological benchmarks. First, the baseline muscle geometries derived from the Visible Human dataset are consistent with MRI-derived population averages. For instance, Handsfield et al.^60^ reported mean volumes of 206.5 cm^3^ for the BFLH, 186.0 cm^3^ for the ST, and 135.2 cm^3^ for the TA. These values compare favorably with our model’s initial volumes of 252.4 cm^3^, 190.1 cm^3^, and 132.4 cm^3^, respectively, confirming that our realistic meshes are representative of healthy adult anatomy. Regarding the link between mechanical workload and biological response, our model successfully predicted the dose-dependent nature of hypertrophy. We observed that the accretion of myofibrils is highly sensitive to the exercise regimen, with high-frequency, high-intensity protocols inducing the steepest monotonic increase. This prediction aligns with the systematic review by Schoenfeld et al^61^, which identifies a significant positive relationship between weekly training frequency (in the range 1-3x/week) and muscle growth. Building on this biochemical foundation, the predicted magnitude of macroscopic hypertrophy aligns well with experimental data, particularly regarding total tissue volume. Our simulation of the DeFreitas (MWF) protocol on realistic geometries predicted a total volume increase of nearly 9%. This is in good agreement with Kubo et al^62^, who reported mean muscle volume increases of 10.1-11.3% in the pectoralis major following 10 weeks of training. Interestingly, the associated increase in mid-belly cross-sectional area was more conservative (∼4-6%) than the range observed in comparable resistance training studies. Specifically, our predictions are slightly lower than the 9.6% increase in thigh muscle CSA reported by DeFreitas et al^45^, the 10% increase found by Damas et al.^63^ in vastus lateralis after 10 weeks of resistance training, and the 7.4% increase reported by Seynnes et al^64^ in the quadriceps femoris following a five-week high-intensity resistance training program. This divergence between the robust volumetric growth and moderate CSA growth highlights the influence of realistic 3D geometries. Unlike idealized geometries where volume scales predictably with CSA (Figure 4C,D), anatomical muscles distribute added mass non-uniformly. Thus, our results suggest that single-slice CSA measurements may underrepresent the total hypertrophic response in complex muscle shapes.

While the proposed framework captures key mechanistic links between signaling and tissue mechanics, we note some limitations. First, the signaling model is a reduced representation of the underlying biochemical network and omits additional pathways and fast-timescale dynamics. A more complex signaling model could give a more accurate description of the underlying dynamics, but at the cost of higher computing demands and the introduction of parameters that are hard to estimate^65^. Second, we model growth strictly as transverse expansion (increasing CSA), driven directly by signaling output. In reality, physiological growth is a response to local mechanical stimuli, such as tissue stress or strain, which trigger mechanotransduction pathways. Furthermore, our model does not account for along-fiber remodeling, known as sarcomerogenesis^66^. This means the model captures how the tissue thickens (hypertrophy) but misses how fibers lengthen to adapt to chronic stretch, an important aspect of tissue remodeling. Crucially, the distinction between these two mechanisms relies entirely on the underlying fiber architecture, as local orientation dictates the axis of both force generation and tissue expansion. While we used Stokes-flow estimates to approximate fiber paths, these are mathematical idealizations. Where available, subject-specific fiber fields derived from diffusion tensor imaging (DTI) could provide improved physiological accuracy compared to the Stokes-flow estimates used here^51^. Nevertheless, this cohort serves as a valuable testing ground to demonstrate the model’s sensitivity to geometry and its potential for personalized simulations.

This study establishes a modular computational infrastructure for linking cellular biology to organ-level morphology. A key feature of this mechanochemical framework is its extensibility. The reduced IGF1-AKT model serves as an adaptable component that can be substituted with more expansive systems biology models to capture phenomena such as metabolic fatigue, calcium handling, or gene regulation. Similarly, the kinematic description of growth can be extended to include along-fiber remodeling (sarcomerogenesis)^67^. Incorporating this mechanism in future iterations could enable the differentiation between concentric-induced thickening and eccentric-induced fascicle lengthening, an important capability for analyzing diverse training modalities. Beyond mathematical extensions, our framework lays the groundwork for integrating the next generation of complex anatomical data. Now that the capability to model growth on realistic geometries is established, there is a need for larger geometric cohorts and subject-specific fiber architectures derived from DTI. Our platform is ready to incorporate such datasets as they become available, allowing for the rigorous characterization of how anatomical variability dictates functional adaptation. Ultimately, by bridging the gap between molecular drivers and macroscopic function, our computational framework lays the groundwork for simulating individual-specific scenarios, from optimizing athlete peak performance to designing targeted rehabilitation protocols for muscle atrophy.

## Acknowledgments

The authors are grateful to Henrik Finsberg and Jørgen Dokken for their assistance with the FEniCSx finite element implementation, and to Halvor Herlyng for generously sharing his implementation of Stokes flow with slip boundary conditions. This work was supported in part by the Wu Tsai Human Performance Alliance at the University of California, San Diego (to P.R.). I.S.D is supported by the Simula–UCSD–University of Oslo Research and PhD training (SUURPh) program, an international collaboration in computational biology and medicine funded by the Norwegian Ministry of Education and Research. M.E.R. acknowledges support from Stiftelsen Kristian Gerhard Jebsen via the K. G. Jebsen Centre for Brain Fluid Research, Wellcome via Award 313298/Z/24/Z (Next-generation simulation and learning in imaging-based biomedicine), and from the Research Council of Norway (RCN). Simulations were performed on the Experimental Infrastructure for Exploration of Exascale Computing (eX3), which is financially supported by the RCN under contract 270053.

## Author contributions statement

I.S.D., M.E.R., and P.R. conceived and designed the project. I.S.D. developed the code and conducted the experiments. I.S.D., M.E.R., and P.R. analyzed the results. I.S.D. prepared the figures. I.S.D., M.E.R., and P.R. wrote the manuscript. All authors edited and reviewed the manuscript.

