## Supplementary Information for "Mechanochemical modeling of exercise-induced skeletal muscle hypertrophy"

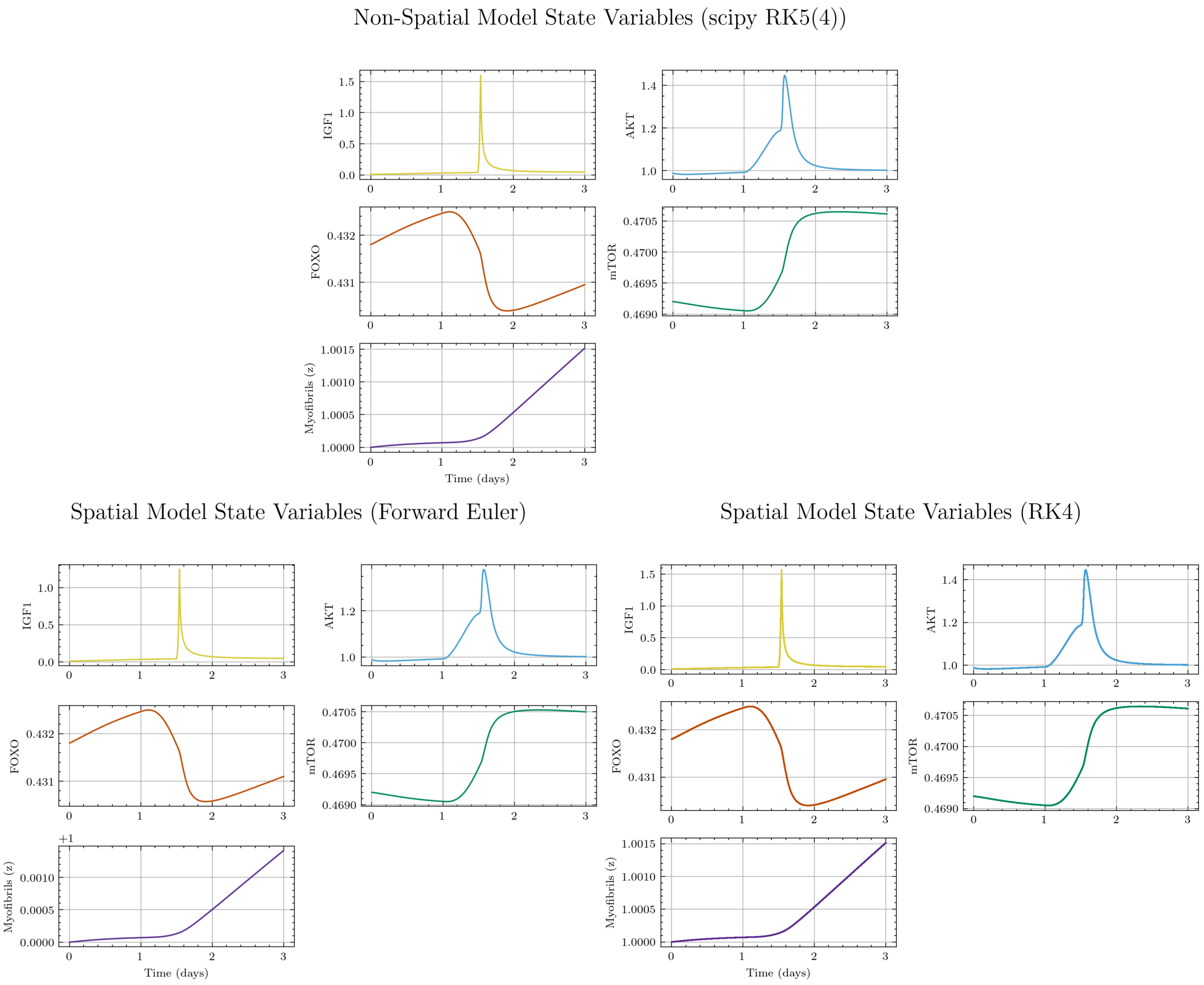


**Supplementary Figure 1. ODE solver verification.** Time course of the ODE state variables comparing the non-spatial SciPy RK45 reference and spatial integrators (forward Euler, RK4) for a single exercise session with homogeneous parameters and time step $\Delta t = 0.05$ h. The spatial RK4 closely matches the RK45 solution.

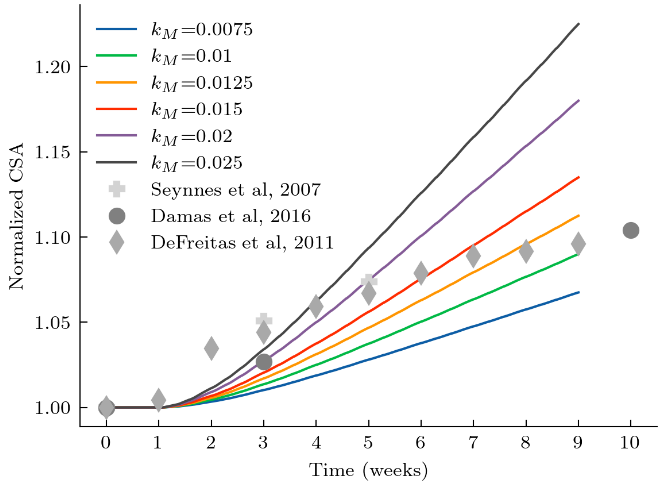


**Supplementary Figure 2. Parameter fit of the protein synthesis rate $k_M$.** To verify the model's physiological relevance, we compared the simulated growth against experimental cross-sectional area (CSA) data. The plot shows the cross-sectional area (CSA) over time for different values of the protein synthesis rate $k_M$, for the DeFreitas (MWF) exercise protocol. The results are compared to experimental data from DeFreitas et al.[1] (diamonds), Seynnes et al.[2] (plus), and Damas et al.[3] (circles). The chosen value of $k_M = 0.01\,\mathrm{h}^{-1}$ (green line) provides a good fit with the experimental data. Simulations were run on a cylindrical geometry with no feedback.

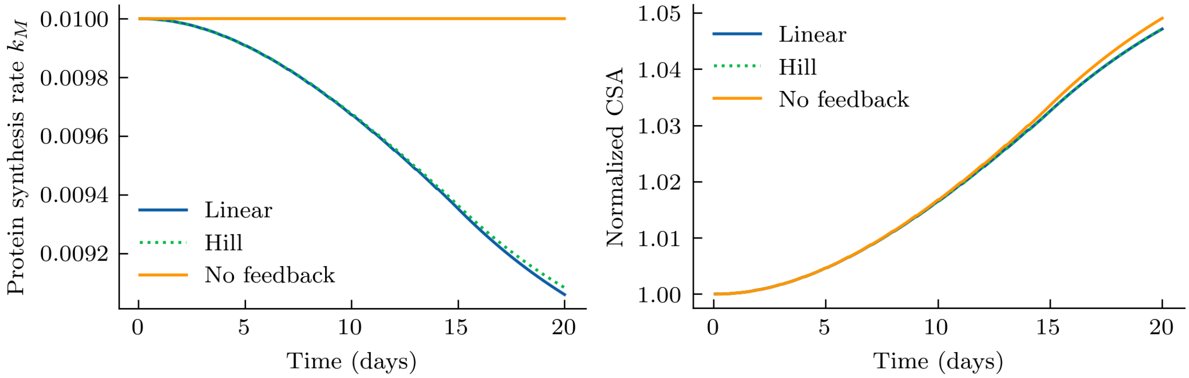


**Supplementary Figure 3. Comparison of Hill-type, linear, and no feedback.** The left panel shows the evolution of the protein synthesis rate, $k_M$, resulting from feedback based on the cross-sectional area, which is shown in the right panel. The simulations were performed on a cylindrical geometry using an exercise protocol of 1 hour of daily exercise for 15 days followed by 5 days of rest. Linear feedback was defined as $k_M(t) = (-2 \cdot A_{CS}(t)/A_{CS}(0) + 3)k_M^0$.

**Supplementary Algorithm 1. Cumulative growth implementation.** Pseudocode for the cumulative growth simulation combining the signaling ODE model with the hyperelastic finite element model.

```
Initialize G_k, G^(k), A^(k), F^(k) to identity tensors.
for (exercise, growth period) pair do
    Simulate ODEs for exercise and subsequent growth period.
    Divide growth period in steps of size Δt.
    for growth step k in growth period do
        Construct incremental growth tensor G_k from ODE solution.
        Update total growth tensor G^(k) = (A^(k-1))^(-1) · G_k · A^(k-1) · G^(k-1).
        Solve hyperelastic model for F^(k) = A^(k) G^(k).
        Update total elastic tensor A^(k) = F^(k) (G^(k))^(-1).
        Compute current cross-sectional area A_CS(t).
        Update protein synthesis rate k_M through feedback function.
    end for
end for
```
